# Batch Production of High-Quality Graphene Grids for Cryo-EM: Cryo-EM Structure of *Methylococcus capsulatus* Soluble Methane Monooxygenase Hydroxylase

**DOI:** 10.1101/2021.12.26.474209

**Authors:** Eungjin Ahn, Byungchul Kim, Uhn-Soo Cho

## Abstract

Cryogenic electron microscopy (cryo-EM) has become a widely used tool for determining protein structure. Despite recent advances in instruments and algorithms, sample preparation remains a major bottleneck for several reasons, including protein denaturation at the air/water interface and the presence of preferred orientations and nonuniform ice layers. Graphene, a two-dimensional allotrope of carbon consisting of a single atomic layer, has recently attracted attention as a near-ideal support film for cryo-EM that can overcome these challenges because of its superior properties, including mechanical strength and electrical conductivity. Graphene minimizes background noise and provides a stable platform for specimens under a high-voltage electron beam and cryogenic conditions. Here, we introduce a reliable, easily implemented, and reproducible method of producing 36 graphene-coated grids at once within 1.5 days. The quality of the graphene grids was assessed using various tools such as scanning EM, Raman spectroscopy, and atomic force microscopy. To demonstrate their practical application, we determined the cryo-EM structure of *Methylococcus capsulatus* soluble methane monooxygenase hydroxylase (sMMOH) at resolutions of 2.9 and 2.4 Å using Quantifoil and graphene-coated grids, respectively. We found that the graphene-coated grid has several advantages; for example, it requires less protein, enables easy control of the ice thickness, and prevents protein denaturation at the air/water interface. By comparing the cryo-EM structure of sMMOH with its crystal structure, we revealed subtle yet significant geometrical differences at the non-heme di-iron center, which may better indicate the active site configuration of sMMOH in the resting/oxidized state.

**Table of Contents (Graphical abstract):** 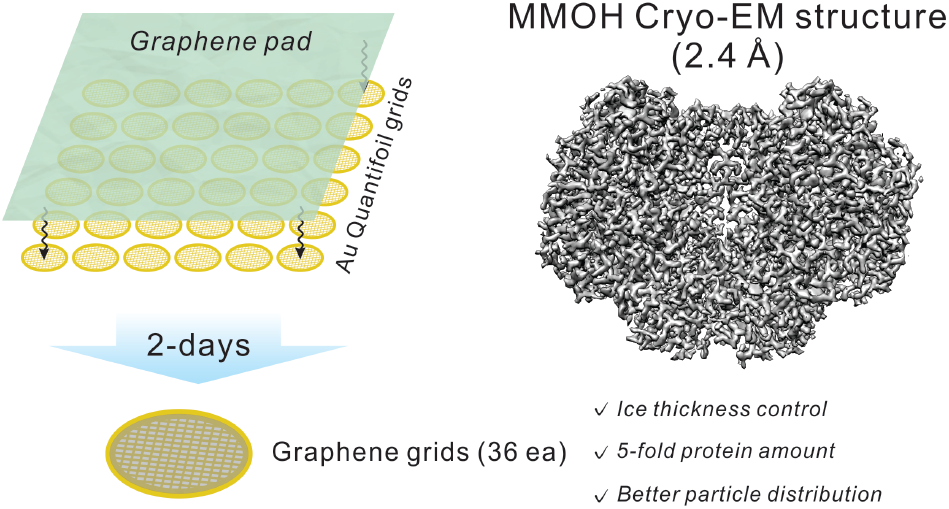

## INTRODUCTION

Single-particle cryogenic electron microscopy (cryo-EM) has evolved as a major technique to determine high-resolution protein structure. Advances in multidisciplinary technologies including direct electron detectors, energy filter systems, advanced algorithms, and data collection/processing strategies have enabled cryo-EM to become a versatile and routine method for the structure determination of biomacromolecules.^1–6^

With these advances and the broad adoption of cryo-EM techniques in instrument development and software algorithms, specimen preparation and grid development have become increasingly important for determining protein structures by cryo-EM.^7–9^ Several challenges remain in cryo-EM sample preparation, such as protein denaturation mediated by the air/water interface (AWI),^10,11^ nonuniform ice thickness, preferred particle distributions/orientations,^12^ and beam-induced motion.^13–15^ To overcome these challenges, the addition of a continuous thin layer of supporting film on the cryo-EM grid has been widely considered and tested; film materials include inorganic metal alloys (e.g., titanium–silicon and nickel–titanium), carbon nanomembranes, and other forms of amorphous carbon.^16–20^ However, these films typically add significant background noise to the microscopic images. Other types of supporting films, such as a lipid monolayer^21^ and two-dimensional (2D) streptavidin crystals,^22^ have been successfully employed to determine protein structures;^23–25^ however, technical challenges have limited their availability and applicability.

An ideal supporting film for cryo-EM grids must have several properties. First, the material must be thin and sufficiently transparent that it does not interrupt the electron beam pathway to minimize unwanted scattering. Second, it should be physically strong enough to stably hold both a thin ice layer and particles during screening and data collection. Finally, it should be electrically conductive to prevent charge accumulation on the surface during lengthy automated data collection. Graphene, a 2D single atomic carbon layer, meets most of the conditions for use as a supporting film, as it possesses electrical conductivity (~15,000 cm^2^·V^−1^·s^−1^), optical transparency (~97.7%), mechanical strength (~1,000 GPa), and minimal scattering events under a 300 kV electron beam.^26–30^ A plasma-treated graphene grid has been shown to achieve more evenly distributed particles in ice and minimize beam-induced particle motion.^31^

Several research groups recently attempted to coat EM grids with graphene or its derivatives to exploit those superior properties for cryo-EM observation.^10,32–39^ Although these earlier attempts were successful, the methods they used are not easily adopted at the laboratory level for several reasons. 1) Some of the reported methods require specific professional instruments, such as chemical vapor deposition (CVD) equipment for graphene synthesis and a Langmuir–Blodgett trough for uniform graphene oxide coating.^39,40^ In addition, it is necessary to collaborate with an expert in graphene or its derivatives during its synthesis, transfer, and modification. 2) Only a few established methods are available for assessing the quality of graphene-coated grids.^34,36,41^

Here, we report a versatile and easy fabrication method for developing graphene-coated grids. We focused on developing a robust graphene-coating protocol for grids that is reproducible and easy to follow. Using a new method, we routinely prepared 36 graphene-coated grids at once in ~1.5 days. We also focused on establishing proper validation methods to evaluate the success of the graphene-coating process on the grid. The method described here does not require expensive instruments, unique skills, or collaborators in material chemistry.

To test the applicability and quality of the graphene-coated grids, we determined the cryo-EM structures of *Methylococcus capsulatus* (*M. capsulatus*) soluble methane monooxygenase hydroxylase (sMMOH) using the newly developed graphene-coated grid. Methanotrophic bacteria in ambient environments use methane as their sole carbon source by converting methane to methanol. The critical enzymes participating in this conversion are two types of methane monooxygenases; particulate methane monooxygenase (pMMO) and soluble methane monooxygenase (sMMO). In particular, sMMO participates in the following reaction:

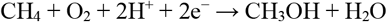

The effectiveness of catalysis by sMMO depends on the interplay of four protein components: sMMO hydroxylase (sMMOH), sMMO reductase (sMMOR), sMMO regulatory subunit (sMMOB), and sMMO inhibitory subunit (sMMOD). The crystal structure of sMMOH, exhibits a 251 kDa heterohexameric (α_2_β_2_γ_2_) architecture with a glutamate- and histidine-coordinated di-iron active site in each α protomer.^42^ sMMOR shuttles electrons from reduced nicotinamide adenine dinucleotide (NADH) to the di-iron center of sMMOH.^43^ sMMOB is an auxiliary component that modulates the reduction potentials at the di-iron center and facilitates substrate availability at the active site.^44,45^ Another regulatory component, sMMOD, inhibits the catalytic activity of sMMOH.^46,47^ To date, all 28 structures of sMMOH deposited since 1993 have been determined solely by X-ray crystallography.^42,48–50^ However, X-ray structures can be affected by high precipitant concentrations and crystal packing. To obtain the atomic-resolution structure without unnatural compounds and crystal contacts, we determined the cryo-EM structure of *M. capsulatus* sMMOH at resolutions of 2.9 and 2.4 Å using Au Quantifoil and the newly developed graphene grid, respectively.

## EXPERIMENTAL DETAILS

### Protocol for graphene transfer to cryo-EM grid

A 25.4 mm × 25.4 mm poly(methyl methacrylate) (PMMA)/graphene pad (Trivial Transfer Graphene) was purchased from ACS Material and stored in a 2–8 °C refrigerator before use. The graphene transfer process was based on the user instructions with several modifications. First, 36 gold Quantifoil holey carbon grids (Au Quantifoil grids) were placed on a three-dimensional (3D)-printed grid transfer tool (Figure S1) immersed in deionized water in a petri dish, and the PMMA/graphene/support textile pad was gently placed on the water surface to prevent water overflow onto the graphene layer. Direct water overflow onto the graphene layer will cause it to shrink, making it unusable. After the water fully permeated the interface between the graphene layer and textile pad (approximately 30 min), the supporting textile was gently pushed to the bottom of the petri dish using tweezers. This procedure separated the graphene layer from the textile pad and caused the graphene layer to float on the water surface (Figure 1). Then the PMMA/graphene layer was carefully matched with the top of the 36 grids using tweezers, and the structure was slowly removed from the water by lifting the grid transfer tool. Residual water in the grid transfer tool was drained with a cleaning wipe, and the PMMA/graphene/grid structure was placed in an oven (100 °C, 30 min) to dry completely. This step also increased the contact between the graphene layer and grids. Fully dried PMMA/graphene-coated grids were individually detached from the grid transfer tool and immersed in acetone solvent (50 °C, 30 min, repeated three times with mild stirring) in a petri dish to dissolve and eliminate the PMMA layer. The acetone-free, air-dried, PMMA-free graphene grids were transferred to a microscope slide glass and baked in the oven (200 °C, 24 h) to evaporate the remaining solvent and swell the polymers on the graphene. To prevent the direct exposure of the graphene grids to heat, we placed the slide glass in a petri dish (glass, with a lid) and also covered it with a glass beaker during baking. The generated graphene grids were individually transferred to a grid box, which was held in a desiccator for up to several months before use.

**Figure 1.**
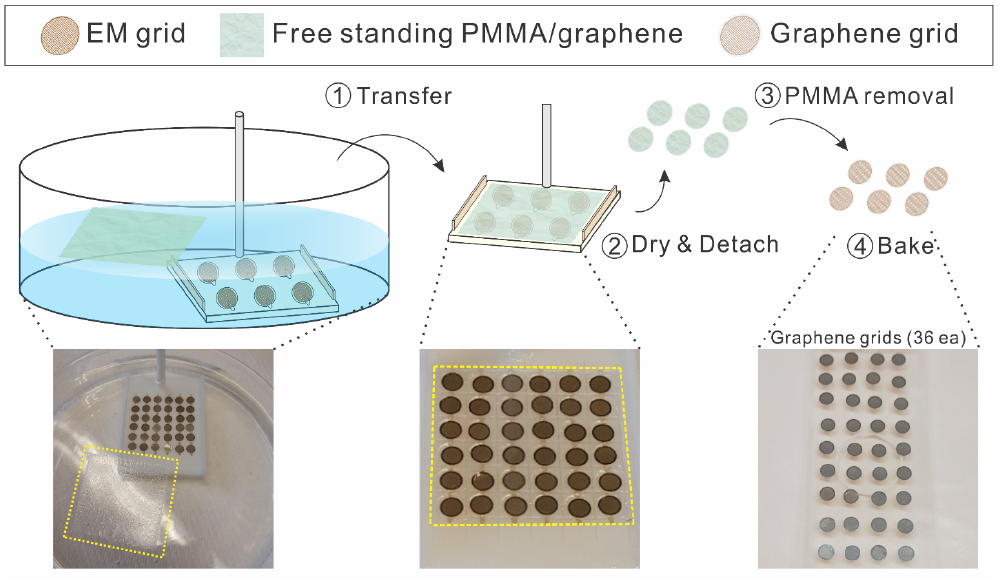
Schematic and photographic views of graphene grid production. Yellow dash square in photographs denotes the PMMA/graphene region.

### Characterizations of graphene grids

Scanning EM (SEM) images were obtained using a NOVA NanoLab 200 instrument with an electron beam power of 5 kV at 0.40 nA. The graphene and other grids were placed on a grid-holding accessory for SEM observation. Transmission EM (TEM) images were obtained at 200 kV using a JEOL-2010F instrument. Raman spectra were collected using a Renishaw instrument equipped with a 532 nm diode laser having a power of 0.5 W; a 1200 lines/mm grating and an Olympus SLMPlan 20× objective lens were also used. All spectra were obtained in the Raman shift range of 1000–3000 cm^−1^ to analyze the framework bands and at peak positions of 2680 cm^−1^ to analyze the 2D band and 1580 and 1380 cm^−1^ to monitor the G and D peaks of graphene (amorphous carbon bands), respectively. The laser was calibrated in static scan mode using a silicon standard. Atomic force microscopy (AFM) images were taken using a Veeco Dimension Icon atomic force microscope with a ScanAsyst-Air AFM tip from Bruker Nano Inc. The sample grids were held on a slide glass by copper tape/slide glass during scanning. The data were analyzed using Nanoscope Analysis 2.0 software. The AFM tip was calibrated in thermal tune mode with a spring constant of 0.4 N/m.

### Purification of *M. capsulatus* sMMOH

*M. capsulatus* (Bath) sMMOH was purified as described in the literature.^45^ In brief, *M. capsulatus* (Bath) was cultured in a nitrate mineral salt medium at 30 °C. Then the harvested cells were collected by centrifugation (11,300*g*) for 20 min at 4 °C. The cell pellet suspended in the buffer was lysed at 4 °C (CV334, Sonics), and the lysate was centrifuged at 30,000*g* for 45 min at 4 °C. The supernatant was collected and filtered through a 0.22 μm membrane. The filtrate was loaded onto DEAE Sepharose, Superdex 200, and finally Q Sepharose columns to obtain sMMOH with >95%purity.

### Cryo-EM sample preparation on graphene grids

Before sample loading, the graphene grids were treated by glow discharge (PELCO easiGlow, Ted Pella Inc.) at 5 mA for 60 s in vacuum (<0.26 mbar). Cryo-EM sample grids were prepared in a Vitrobot Mark IV instrument (Thermo Fisher Scientific), which was set to 4 °C at 100% humidity. Purified protein solution (3 μL) was loaded onto the graphene surface of the graphene grid or holey carbon surface of the Au Quantifoil grids as indicated. After 30 s, the grids were blotted for 4 s with a blot force of 0 and immediately plunged into precooled liquid ethane for vitrification.

### Data acquisition

A total of 3880 (graphene grid) and 3075 (Au Quantifoil grid) raw movie stacks were automatically collected on a 300 kV Titan Krios using a K3 direct electron detector (with a BioQuantum energy filter, Gatan) at the Pacific Northwest Center for Cryo-EM (PNCC). Raw movies from Quantifoil and graphene grid were collected in K3 super-resolution mode at a magnification of 81,000× (slit width 20 eV, spot size 5, C2 aperture 50 μm) with a super-resolution pixel size of 1,027 and 1.059 Å, respectively. The total exposure time was 3.4 s at 0.85 e^−^/Å^2^ per frame to generate 60-frame gain-normalized microbial reverse-electrodialysis cell stacks. The total dose per stack were 49 and 51 e^−^/Å^2^ for Quantifoil and graphene grid, respectively^2^.

### Cryo-EM data processing and model refinement

All processing was completed in RELION and cryoSPARC.^2,51^ The MotionCor2 implemented in RELION corrected the initial drift and beam-induced motion, and Gctf was used to measure the contrast transfer function (CTF).^52^ Following CTF estimation, micrographs were manually inspected, and those with outliers in the defocus value, ice thickness, or astigmatism, as well as micrographs with lower predicted CTF-correlated resolution (>5 Å), were removed from further processing. The initial set of particles was selected using Topaz.^53^ The selected particles were further 2D- and 3D-classified by iterative classification and selection rounds using an *ab initio* 3D reconstructed model as a starting reference in cryoSPARC. A total of 325k and 479k particles were chosen to build final 3D reconstruction maps with a resolution of 2.31 and 2.60 Å (FSC_0.143_) for the graphene grid and Au Quantifoil grid, respectively. Using the crystal structure of sMMOH (PDB ID: 1MTY),^54^ we refined the structures using real-space refinement in the PHENIX program and COOT.^55,56^ The reported resolutions of the final maps (2.4 Å for the graphene grid and 2.9 Å for the Au Quantifoil grid) were estimated using the map versus model FSC curves (FSC_0.5_^map vs. model^) in phenix.xtriage.^55^ UCSF Chimera^57^ was used to visualize the EM density and obtain illustrations for figures. The structural data has been deposited to the PDB and EMDB. The accession numbers of *M. caps* MMOH structures for the graphene grid and Au Quantifoil grid are PDB: 7TC8 / EMDB-25805 and PDB: 7TC7 / EMDB-25804, respectively.

## RESULTS AND DISCUSSION

### Making graphene-coated Au Quantifoil grids

The graphene transfer approach we developed and adopted is based on the polymer-film-assisted transfer method instead of the polymer-free transfer method.^58–60^ Although the polymer-free transfer method has several advantages (for example, it can produce a hyperclean graphene surface and graphene with high-order crystallinity), skills and knowledge are required for interfacial control between the transfer solvents, and thus the method is not suitable for those who are unfamiliar with the handling of graphene and surface engineering. By contrast, polymer-assisted graphene transfer has several advantages that make it possible to overcome these challenges. 1) It provides an intuitive and easy-to-use method of generating graphene-coated Au Quantifoil grids (i.e., graphene grids) at the laboratory level for those who are unfamiliar with nanomaterials. 2) It can be used for the mass production of high-quality graphene grids with good surface coverage (>95%) and cleanness. It takes 1.5 days to generate a batch of 36 graphene grids using the method we developed. The method requires a petri dish, an oven, a 25.4 mm × 25.4 mm PMMA/graphene pad (ACS Material), acetone, and a 3D-printed graphene transfer tool that we designed (Figure S1). The graphene grid synthesis process is described in detail in the Experimental Details section and illustrated in Figure 1. In particular we note that the baking step (200 °C, overnight) in the oven was crucial for producing graphene grids with clean surfaces (Figure S2). The boiling point of the PMMA rinsing solvent (i.e., acetone) is 56 °C. However, we found that a moderate baking temperature (100 °C) was not sufficient to remove the swelled solvent in the PMMA residues (Figure S2a). We recommend performing the baking step without vacuum assistance, as oxygen species facilitate the removal of PMMA residues during this process (Figure S2b). By increasing the temperature to 200 °C, we minimized the residual PMMA on the graphene surface without damaging the coated graphene (Figure S2c).^61^ Although residual PMMA on the graphene surface is inevitable because it adheres to surface defects/boundaries during graphene synthesis by CVD, it can be minimized by the high-temperature baking process.^62^ We also noticed that two layers of protection (the lid of the glass petri dish and the beaker) during overnight baking prevent direct exposure to heat and thus minimize the damage to the graphene and the holey carbon. Both copper and gold Quantifoil grids were examined to test which grid material is more suitable for the graphene coating procedure (Figure S3). We found that the Au Quantifoil grid exhibits better graphene coating because the Cu Quantifoil grid become heavily oxidized during baking, as indicated by a color change and more damage to the graphene/Quantifoil holey carbon during overnight baking at 200 °C.

### Characterization of the graphene grid

Previous studies in generating graphene-/graphene-oxide-coated grids have demonstrated the coating quality by testing and visualizing standard macromolecules such as the 30S ribosomal subunit^36^ and apo-ferritin^37^ using cryo-EM.^32^ Although this approach may indirectly indicate the quality of graphene or graphene-oxide grids, some or all of the cryo-EM data must be used to examine the quality of the grids, which is time-consuming and poses a potential risk to the precious specimen on the grid. Therefore, easily applied quality control/validation tools are urgently needed before sample application. As quality control tools for the graphene grids, we employed four approaches: SEM, bright-field (BF) TEM, Raman spectroscopy, and AFM.

The graphene-coated Au Quantifoil grids were characterized using SEM and BF TEM to examine the coverage and surface quality. First, we compared the untreated PMMA/graphene-coated Au Quantifoil grids (before PMMA removal) and PMMA-free graphene grids. As shown in Figure 2d, the surface of the PMMA/graphene grid was fully covered with a thick PMMA layer (approximately 500 nm according to the product information), which even blocked the observation of the hole patterns in the commercial Quantifoil grid (Au 300 mesh, R1.2/1.3). The TEM image and selected area electron diffraction (SAED) pattern of the PMMA/graphene grid (Figure 2e,f) showed a strong scattering pattern of amorphous carbon, which originated from the covered PMMA layer, and the characteristic SAED pattern of monolayer graphene (Figure 2f).^63,64^ After PMMA removal by rinsing and baking, most of the thick PMMA layer was eliminated from the graphene surface, as confirmed by SEM, BF TEM, and SAED observations (Figure 2g–i). The SEM image indicates that the graphene fully covers the hole-patterned carbon of the Au Quantifoil grid. The SAED diffraction image further demonstrates that the diffraction pattern of the amorphous carbon ring is minimized, except for residual signals originating from the supporting holey carbon film of the Quantifoil grid. The graphene transfer method seems to be applicable to various other types of EM grids, but the Au Quantifoil showed the best graphene coverage (>95%) and cleanness. We note that materials with larger mesh number and smaller hole diameter showed better graphene coverage and less damage to the graphene grids during baking. Smaller hole diameters have also been shown to minimize beam-induced particle motion.^9^

**Figure 2.**
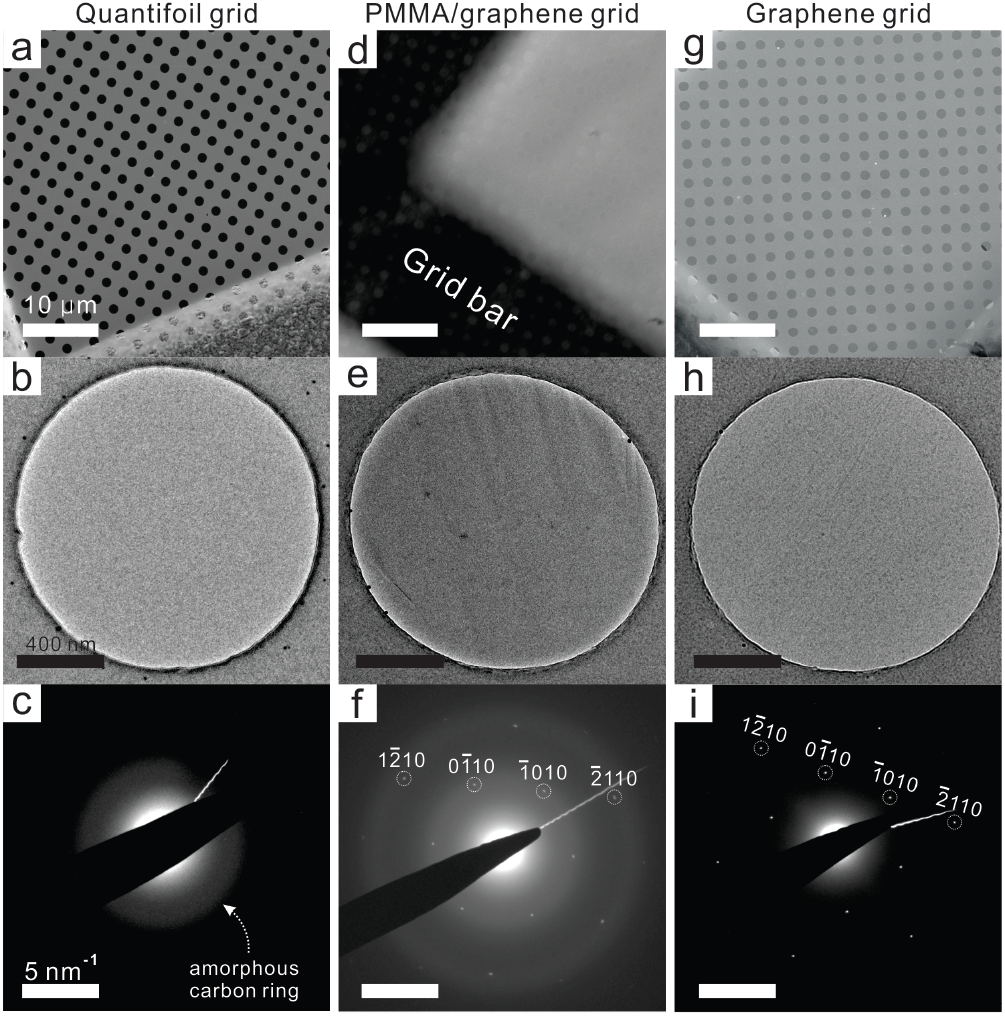
(a-c) Left-side images are from untreated commercial Au Quantifoil grids. (d-f) Mid-side images are from PMMA/graphene-coated Au Quantifoil grids. (g-i) Right-side images are PMMA-free graphene grids after rinsing/baking steps. (a, d, g) Scale bars in SEM image represent 50 μm. (b, e, h) Scale bars in TEM image represent 400 nm. (c, f, i) Scale bars in SAED images represent 5 nm^−1^.

Figure 3 shows the Raman spectra of commercial monolayer graphene on Cu foil (Graphenea; top), the commercial Quantifoil grid (Electron Microscopy Sciences; middle), and the inhouse graphene-coated grid (bottom). Single-layer graphene without defects was observed in the commercial graphene grid and confirmed by the absence of the D peak (1348 cm^−1^; defect-induced second-order Raman scattering) and the high peak intensity ratio between the G peak (1580 cm^−1^; in-plane vibrations of sp^2^-hybridized carbon atoms) and the 2D peak (2670 cm^−1^; second-order overtone of a different in-plane vibration).^65,66^ It is generally acknowledged that a Raman intensity ratio of the 2D and G peaks (*I*_2D_/*I*_G_) of >2.0 denotes single-layer graphene.^67,68^ The Raman spectra of the Quantifoil grid (before graphene transfer) shows large, broad D and G bands, which are derived from the hole-patterned amorphous carbon.^69^ After graphene transfer to the Quantifoil grid, both the characteristic G and 2D peaks of the graphene layer and the D and G band patterns of the Quantifoil grid appear in the Raman spectrum of the graphene-coated grid, indicating that the graphene monolayer was successfully transferred and coated on the Quantifoil grid.

**Figure 3.**
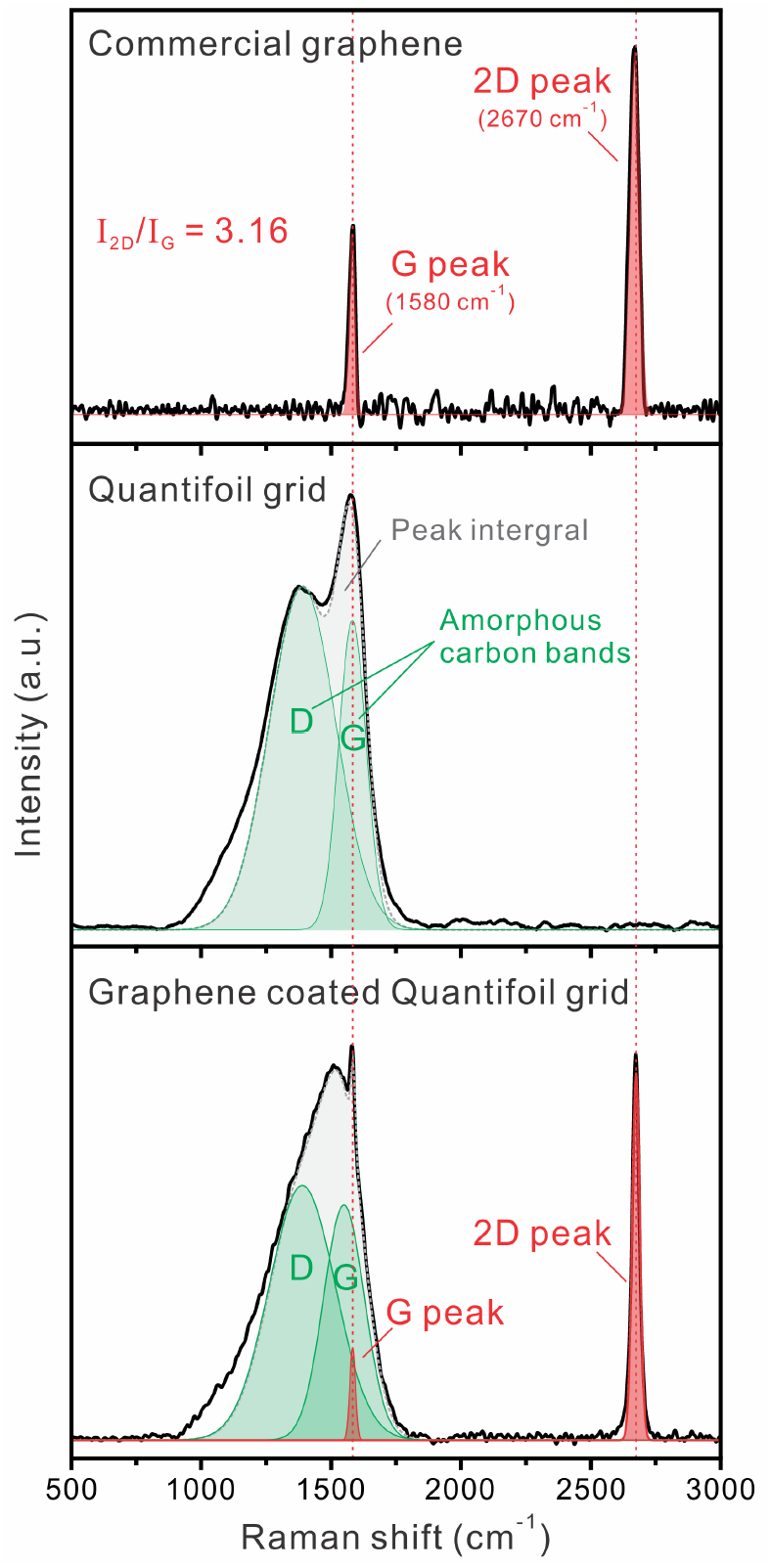
Raman spectra of commercial graphene monolayer (top), commercial Au Quantifoil grid (middle), and graphene-coated Au Quantifoil grid (bottom). Red-dotted vertical line denotes the peak positions (G and 2D) of graphene. Green colored region denotes the Raman band positions (D and G) of amorphous carbon.

AFM was further employed to characterize the morphology and physical properties of the graphene grids. The Quantifoil grid has regularly patterned holes, which were fully covered after the PMMA/graphene transfer process (Figure 4a–d). After the thick PMMA layer was removed, the hole pattern covered by graphene appeared, with minimal surface residues (Figure 4e,f); most of the areas of interest (cryo-sample positions) were clean and defect-free. The graphene grids exhibited a relatively large *z*-scale variation (300 nm), mainly because of a fragile grid wrinkle that occurred during AFM sample preparation. Inside of the graphene-covered holes, the surface was flat and clean, and showed low *z*-scale variation (10 nm) (Figure 4g,h). Therefore, the Raman spectroscopy and AFM results, in addition to SEM and TEM images, can be good validation tools for examining the quality of graphene-coated grids after graphene transfer and before specimen application.

**Figure 4.**
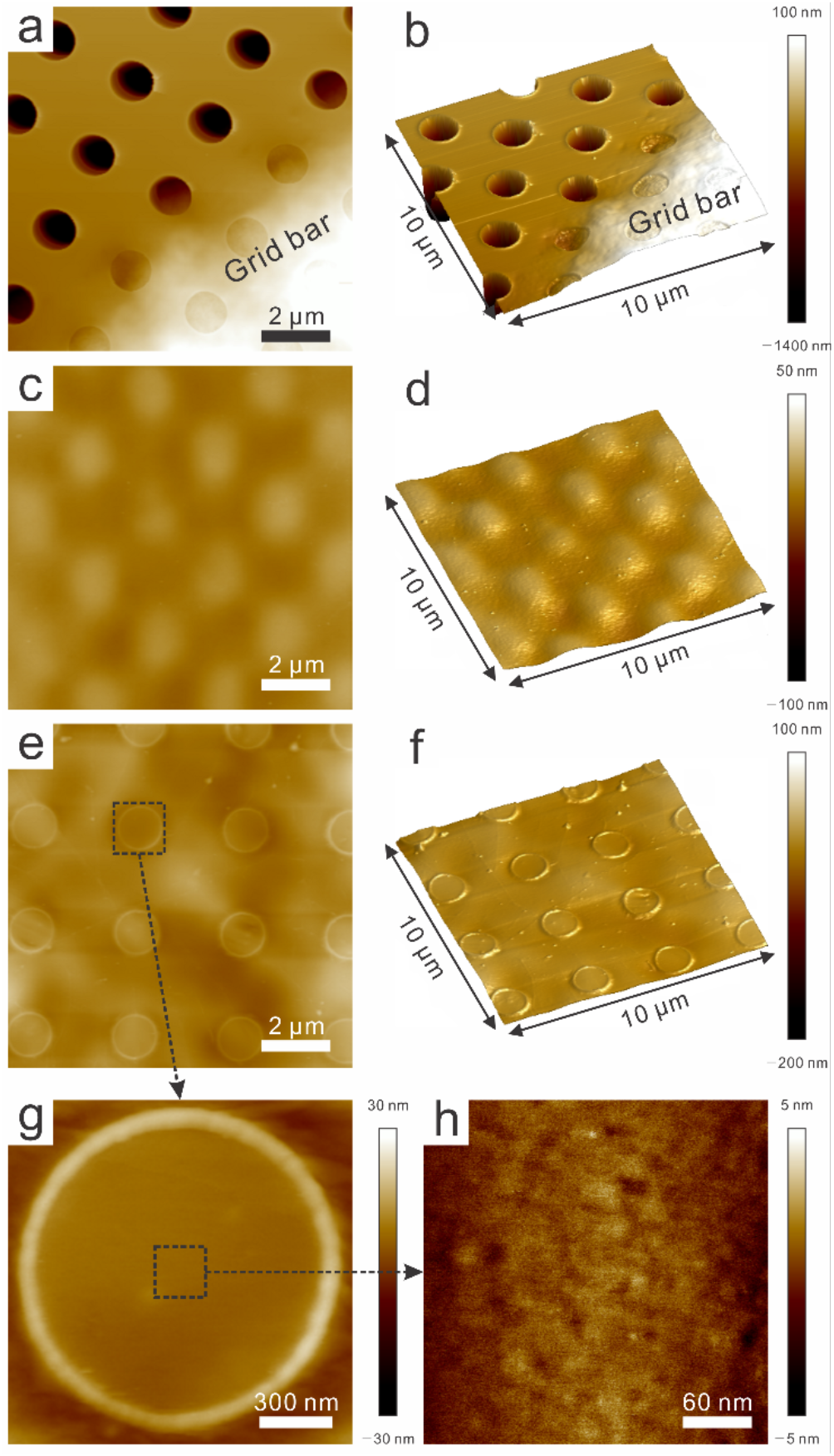
AFM height mode and 3D mode images of the Quantifoil, PMMA/graphene coated Quantifoil, and PMMA-free graphene grids. Note that the *z*-scale is not the same in these images. (a) Height mode and (b) 3D image of Quantifoil grid with 1.5 μm z-scale. (c) Height mode and (d) 3D image of PMMA/graphene coated Quantifoil with 150 nm *z*-scale. (e) Height mode and (f) 3D image of PMMA-free graphene grid with 300 nm *z*-scale. Magnified region of black dotted box for (g) single hole and (h) center of the hole with 60 and 10 nm *z*-scale, respectively.

### Cryo-EM structure determination of *M. capsulatus* sMMOH

Since Rosenzweig and coworkers determined the first crystal structure of *M. capsulatus* sMMOH in 1993,^42^ the structures of different enzymatic stages of sMMOH have been determined by X-ray crystallography.^42,70–72^ Earlier studies indicated that the geometry of the non-heme di-iron center plays a critical role in indicating the enzymatic stage of sMMOH. The di-iron geometry, which is coordinated by two histidines (His147 and His246) and four glutamates (Glu114, Glu144, Glu209, and Glu243), undergoes subtle yet significant conformational changes depending on its catalytic reaction stage and its association with auxiliary subunits (e.g., MMOB and MMOD).^45,47^ This rearrangement of the di-iron coordination is interlocked and triggered by the movement of the four-helix bundle (helices B, C, E, and F) that contain these two histidines and four glutamates. Although currently available crystal structures of sMMOH have provided valuable information regarding di-iron coordination shifts under various circumstances, protein crystallization requires high precipitant concentrations and unnatural compounds, and the protein structure can be further constrained by crystal contacts. To illustrate the structure of sMMOH in solution and validate the general applicability of the graphene-coated grid, we determined the cryo-EM structures of *M. capsulatus* sMMOH at resolutions of 2.9 and 2.4 Å using the Au Quantifoil and graphene-coated grids, respectively, under normal buffer conditions [30 mM 4-(2-hydroxyethyl)-1-piperazineëthanesulfonic acid, pH 7.0, 150 mM NaCl, and 1 mM tris(2-carboxylethyl)phosphine) (Figure 5a–c).

**Figure 5.**
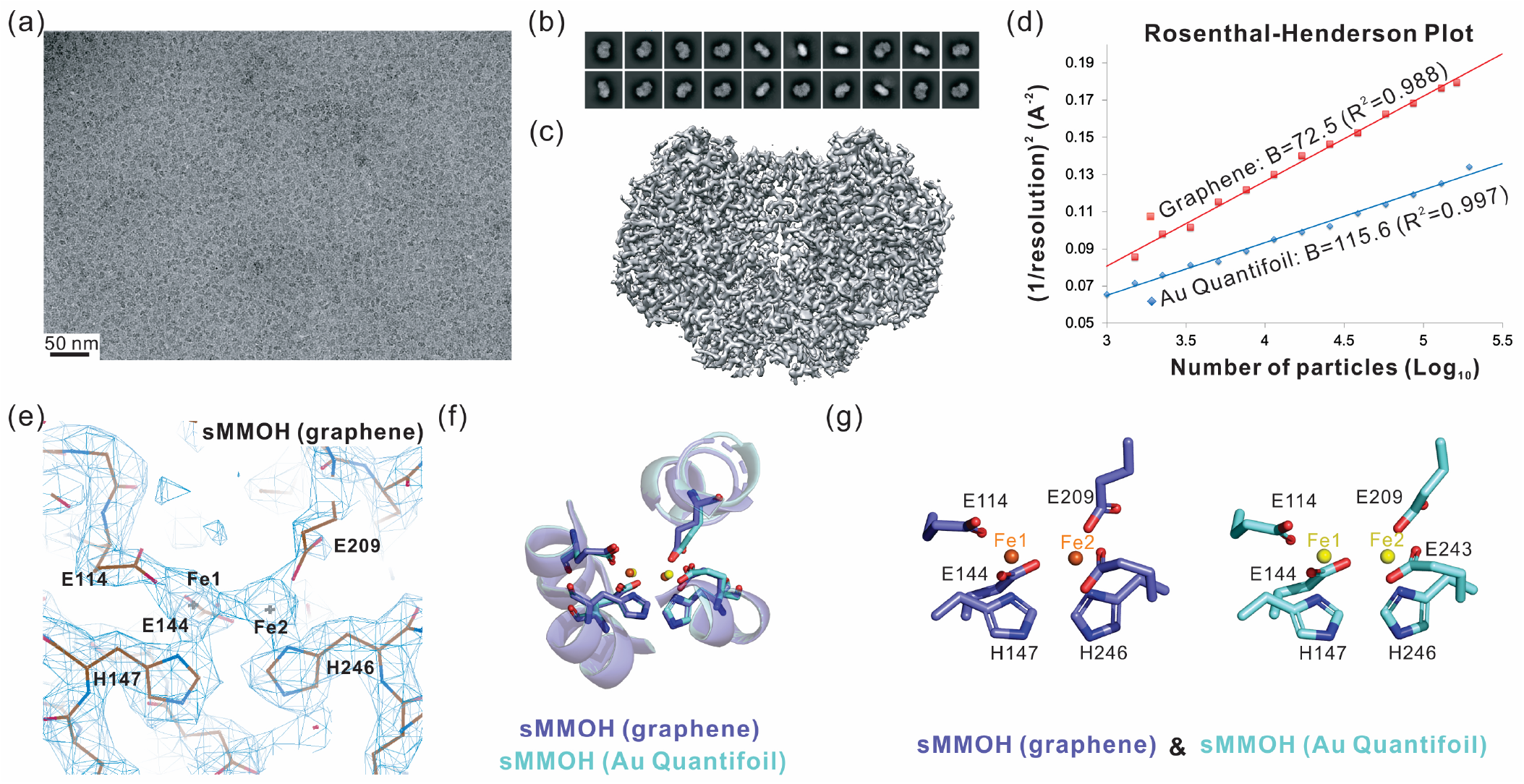
(a) Representative microscopic image of *M. caps* sMMOH in the graphene grid (300 kV). (b) Top 20 2D classes selected from 200 classes of sMMOH in the graphene grid. (c) Reconstructed 3D cryo-EM map (2.4 Å resolution) of sMMOH in the graphene grid. (d) The Rosenthal-Henderson plot of sMMOH structures determined using the Au Quantifoil (blue diamond-shape) and graphene (red square) grids. (e) Cryo-EM map of non-heme di-iron center of sMMOH (graphene grid) coordinated by four Glutamates and two histidines. The image was captured by COOT^56^ (f) Structural overlay of sMMOH four-helix bundle (helix B, C, E, and F) and (g) Coordinating residues (four glutamates and two histidines) determined by Au Quantifoil (cyan; yellow for two irons) and graphene (blue; orange for two irons) grids.

### Structural comparison of sMMOH determined using Au Quantifoil and graphene-coated grids

Purified sMMOH at concentrations of 1.3 and 0.5 mg/mL was applied to the Au Quantifoil and graphene-coated grids, respectively, to prepare them to receive the specimen (Table S1). A total of 3075 (Au Quantifoil grid) and 3880 (graphene grid) raw movie stacks were collected using 300 kV Titan Krios with a K3 direct electron detector equipped with a BioQuantum energy filter (Figures 5a and S4a). Particles at concentrations of 2.1 M (Au Quantifoil grid) and 4.2 M (graphene grid) were obtained after particle selection using the TOPAZ program.^53^ After iterative 2D and 3D classification (Figures 5b and S4b), 479k and 325k particles were selected for the Au Quantifoil grid and graphene grid to reconstitute the cryo-EM structures of sMMOH at resolutions of 2.9 and 2.4 Å, respectively, after real-space refinement by PHENIX^55^ (Table S2, Figures 5c and S4c). The 2.4 Å cryo-EM structure of sMMOH obtained using the graphene grid shows clear side chain densities throughout the entire structure, including the non-heme di-iron center (Figure 5e). The Rosenthal–Henderson plot indicates that the quality and behavior of the final selected particles from the graphene grid are slightly better than those of the particles from the Au Quantifoil grid (Figure 5d). We also noted that the graphene grid requires approximately five times less protein to obtain a similar number of particles (Table S1). Although we found that sMMOH exhibited a preferred orientation on the graphene grid, this orientation might be caused in part by the molecular nature of sMMOH, which has a flat shape (Figures 5b,c and S5). It might be possible to minimize this behavior by using globular particles. It is not clear why the specimen on the graphene grid exhibits better behavior than that on the Au Quantifoil grid, but one possible explanation might be the minimal particle exposure to the AWI. Because particles adhere to the graphene surface after plasma treatment, exposure to the AWI can be dramatically reduced. Russo and Passmore also demonstrated that a plasma-treated graphene grid reduces the beam-induced particle motion, which may further contribute to resolution improvement.^31^ However, the final refined structures of sMMOH on the Au Quantifoil and graphene grids are almost identical, suggesting that structural perturbation by the graphene layer is negligible (Figure S4d). The Cα root-mean-square (rms) difference in the two structures is 0.338 Å, and they have nearly identical di-iron center geometries (Figures 5f,g and S4d).

### Structural comparison of sMMOH determined by cryo-EM and X-rays

To evaluate potential structural perturbation during crystallization, we compared the X-ray and cryo-EM structures of sMMOH. We adopted the X-ray structure of sMMOH in the oxidized/resting state (PDB ID: 1MTY) for comparison.^54^ Purified sMMOH was crystallized under buffer conditions of 25 mM Li_2_SO_4_, 50 mM NH_4_OAc, and 5% polyethylene glycol (PEG 4000), and its structure was determined at a resolution of 1.7 Å. The structural overlay of the crystal structure (PDB ID: 1MTY) and cryo-EM structure (graphene grid) of sMMOH exhibit a Cα rms difference of 0.351 Å, indicating that the architecture of the two structures is very similar overall (Figure S6a). However, we observed differences in the di-iron position and surrounding di-iron coordinating residues (Figure 6). First, we observed differences in the positions of both irons. Fe_1_^3+^ and Fe_2_^3+^ were shifted by 0.5 Å (protomer1)/0.2 Å (protomer2) and 1.1 Å/0.6 Å, respectively, in the cryo-EM structure compared to the X-ray structure (Figure 6).

**Figure 6.**
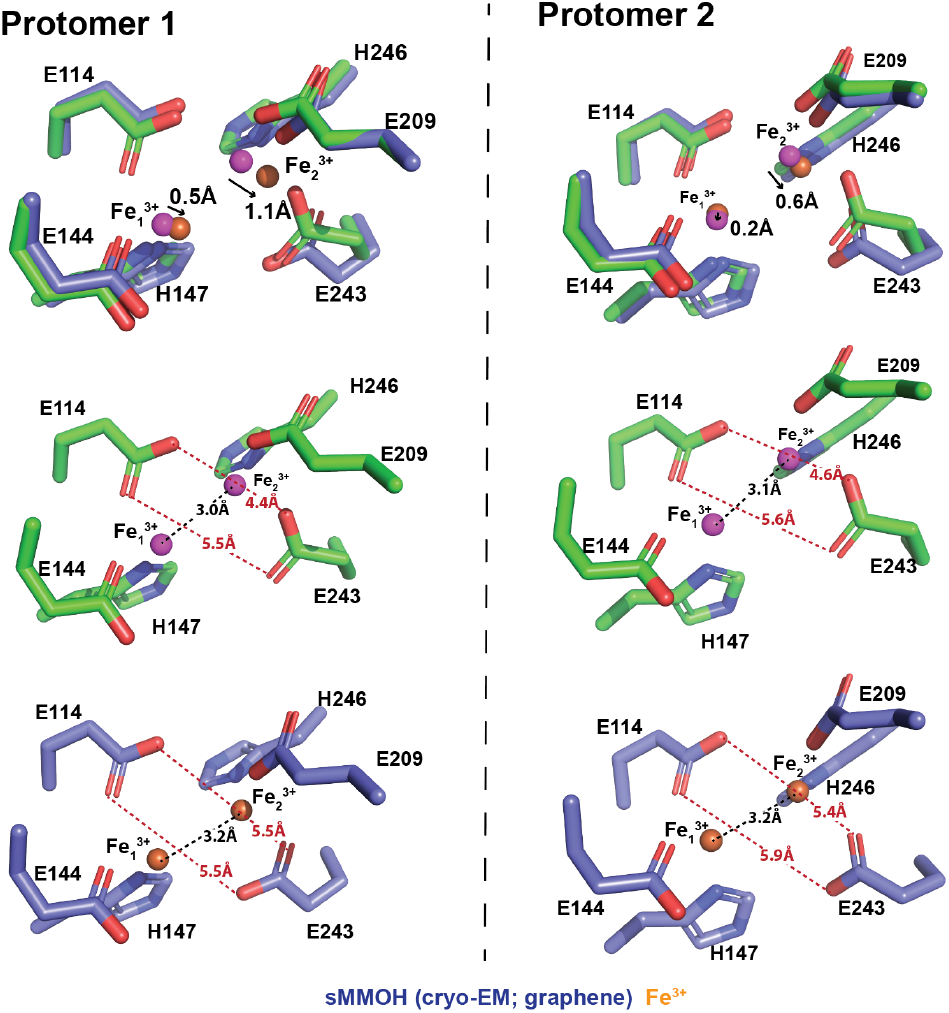
Non-heme di-iron center geometry comparison in between x-ray structure (PDB ID: 1MTY)^54^ and cryo-EM structure (graphene grid). The di-iron center geometry of both sMMOH protomers (protomer 1 and 2) was compared.

The distance between the two γ carboxyl groups of Glu114 and Glu243 also changed, from 5.5 Å/4.4 Å (protomer1) and 5.6 Å/4.6 Å (protomer2) in the X-ray structure to 5.5 Å/5.5 Å (protomer1) and 5.9 Å/5.4 Å (protomer2) in the cryo-EM structure. This result indicates that two γ carboxyl groups of Glu114 and Glu243 moved away from each other in the cryo-EM structure, which resulted in the changes in the positions of the two irons, particularly that of Fe_2_. The iron–iron distances also increased from 3.0 Å (protomer1) and 3.1 Å (protomer2) to 3.2 and 3.2 Å, respectively. We further observed an unidentified cryo-EM density on top of the di-iron center, where the substrates had been located (Figure S7).^47,72,73^ A similar density was also observed in several crystal structures of sMMOH, but one of the crystallization compounds was assigned to refine the structure (e.g., acetate, in PDB ID: 1MMO).^42^ This result indicates that this increase in density might be attributable to one of the sMMOH substrates or molecules involved in the catalytic reaction of sMMOH, which might be co-purified during protein purification. Regardless of the type of compound, it certainly participates in di-iron coordination in the resting state of sMMOH (Figure S7). Because we were not able to identify the nature of this compound, we decided not to build a model of this density.

## CONCLUSION

A versatile method of developing graphene-coated EM grids (i.e., graphene grids) was established using various in situ characterization tools to assess the quality. This method does not require professional skills or expensive instruments and is easily applied to commercially available EM grids with minimal polymeric residues and high coverage. Moreover, high-quality graphene coatings were obtained after cleaning and baking steps. The graphene grids were tested by determining the cryo-EM structure of *M. capsulatus* sMMOH at a resolution of 2.4 Å. We conclude that the graphene grid has advantages for cryo-EM structure determination because it requires five times less protein, can maintain a thin ice layer, achieves evenly distributed particles, and minimizes beam-induced particle motion during data collection. Furthermore, the graphene grid can be applied to fragile proteins, which are prone to denaturation at the AWI.

## ASSOCIATED CONTENT

### Supporting Information

The Supporting Information is available free of charge at https:xxxxxxxx.xxx.

Supporting xxxx (pdf)

## AUTHOR INFORMATION

### Author Contributions

The manuscript was written through contributions of all authors. U.-S.C. designed the research; E.A. and B.K. performed research and analyzed data; E.A., B.K., and U.-S.C. wrote the manuscript. All authors have given approval to the final version of the manuscript.

### Notes

The authors declare no competing financial interest.

## ACKNOWLEDGMENT

This work was supported by grants from R01 DK111465, R01 CA250329, and R01 NS116008 to U.-S.C. A portion of this research was supported by NIH grant U24GM129547, performed at the PNCC at OHSU, and accessed through EMSL (grid.436923.9), a DOE Office of Science User Facility sponsored by the Office of Biological and Environmental Research. We thank Dr. Leila Foroughi and Dr. Adam Matzeger in the Department of Chemistry at the University of Michigan for the use of the Raman spectroscopy instrument and assistance. We thank the staffs at the University of Michigan cryo-EM center for the use of the Morgagni instrument and assistance. We thank Dr. Haiping Sun and the Michigan Center for Materials Characterization for the use of the AFM instrument and assistance.

## ABBREVIATIONS

Cryo-EM: cryogenic electron microscopy
*M. capsulatus* sMMOH: *Methylococcus capsulatus* soluble methane monooxygenase hydroxylase
sMMOR: sMMO reductase
sMMOB: sMMO regulatory subunit
sMMOD: sMMO inhibitory subunit
AWI: air-water interface
CVD: chemical vapor deposition
PMMA: poly(methyl methacrylate)
SEM: scanning electron microscopy
TEM: transmission electron microscopy
AFM: atomic force microscopy
CTF: contrast transfer function
SAED: selected area electron diffraction
rms: root-mean-square

