## Supplementary information for "Batch Production of High-Quality Graphene Grids for Cryo-EM: Cryo-EM Structure of *Methylococcus capsulatus* Soluble Methane Monooxygenase Hydroxylase"

**Table S1.** Comparison of statistics in cryo-EM structure determination (*M. caps* sMMOH)

| | Au Quantifoil grid | Graphene grid |
|---|---|---|
| Protein concentration | 1.3 mg/ml | 0.5 mg/ml |
| Total number of movie stacks | 3075 | 3880 |
| Total number of particles picked | 2.1 M | 4.2 M |
| Total number of particles for the final 3D reconstruction | 479 k | 325 k |
| Resolution ($FSC_{0.143}$) | 2.60 Å | 2.31 Å |
| Resolution ($FSC_{0.5}^{map\ vs.\ model}$) | 2.9 Å | 2.4 Å |

**Table S2.** Cryo-EM data collection, refinement and validation statistics

| | **MMOH on Quantifoil** | **MMOH on Graphene** |
|---|---|---|
| **Data collection** | | |
| Microscope | Titan Krios K3 | Titan Krios K3 |
| Voltage (kV) | 300 | 300 |
| Defocus range (μm) | -0.6 to -1.9 | -0.7 to -2.0 |
| Pixel size (Å) | 1.0275 | 1.059 |
| Total dose ($e^-$ / $Å^2$) | 49 | 51 |
| Particles (initial) | 2.1 mil | 4.2 mil |
| Particles (final) | 479 K | 325 K |
| **Reconstruction** | | |
| Symmetry | C1 | C1 |
| Resolution (unmasked Å) | 3.3 | 2.9 |
| Resolution (mask Å) | 2.60 | 2.31 |
| Map-sharpening *B f*actor ($Å^2$) | 115.6 | 72.5 |
| **Model composition** | | |
| Atoms (Hydrogen) | 20579 (3210) | 20579 (3210) |
| Protein residues | 2113 | 2113 |
| Water | 0 | 0 |
| **Refinement** | | |
| Resolution (Å) | 2.9 | 2.4 |
| *B* factor ($Å^2$) | | |
| Proteins | 34.89 | 13.83 |
| Ligand | 43.47 | 20.74 |
| Model-to-mal fit (CC) | 0.86 | 0.85 |
| **R.m.s. deviations** | | |
| Bond (A) | 0.005 | 0.002 |
| Angles (°) | 0.534 | 0.428 |
| **Validation** | | |
| Clashscore | 14.41 | 16.06 |
| Rotamer outliers (%) | 0 | 2.53 |
| Ramachandran plot (% favored) | 97.67 | 98.33 |
| Ramachandran plot (% outliers) | 0.00 | 0.00 |
| MolProbity score | 1.74 | 2.01 |

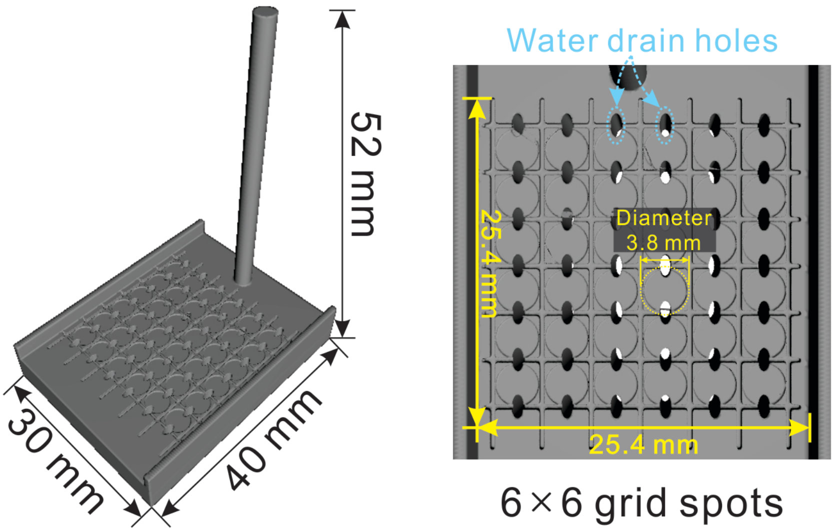


**Figure S1.** Three-dimensional angle view (left) and top view (right) of the 3D-printed grid transfer tool (The Form 3 with the white resin, Formlabs) with scales. The total 36 grid holes (3.8 mm diameter with 0.2 mm depth) are positioned within 25.4 mm × 25.4 mm (1 inch × 1 inch). A groove is positioned in between holes for cutting the graphene layer using tweezers after the drying step (100 °C, 30 min). The graphene grid then can be detached and transferred individually for acetone washing. To hold and position grids during the scooping step in water, we introduced the water drain hole in between grid holes (cyan dot circles). This water drain hole further provides the space to pick the grid (after drying step) without bending it with the tweezers. The STL file for this grid transfer tool can be freely downloaded in the following link. (https://drive.google.com/file/d/1Ypu8qxs3QKvii0EEzsLVLajQxGKETX9k/view?usp=sharing).

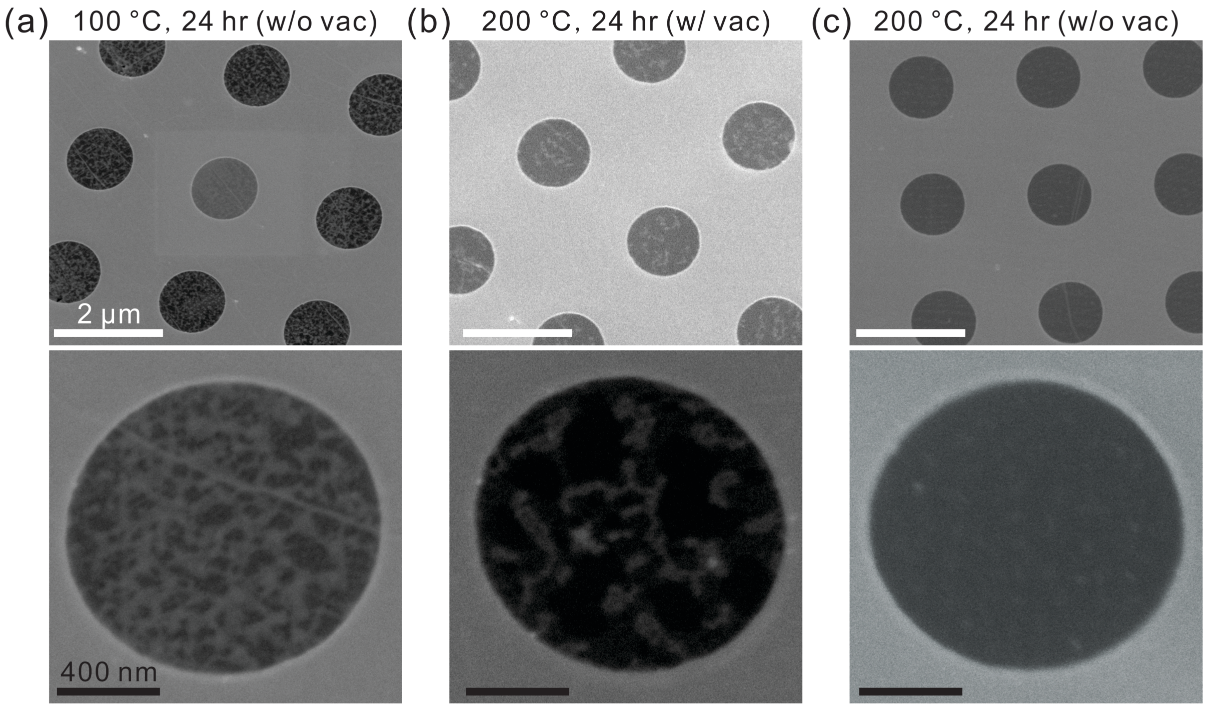


**Figure S2.** SEM images of graphene grids (Au Quantifoil) made in different baking conditions of (a) 100 °C without vacuum, (b) 200 °C with vacuum, and (c) 200 °C without vacuum. Bottom side images are magnified hole images of each condition.

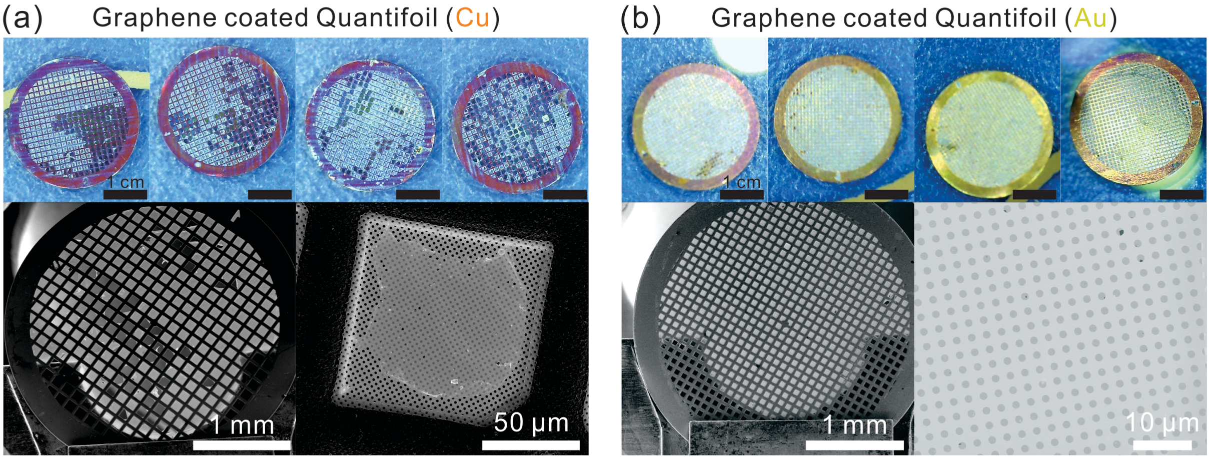


**Figure S3.** Optical microscopy (OM) and SEM images in different magnifications of graphene-coated (a) Cu Quantifoil and (b) Au Quantifoil grids. Note that scale bars in magnified SEM image are in different scale.

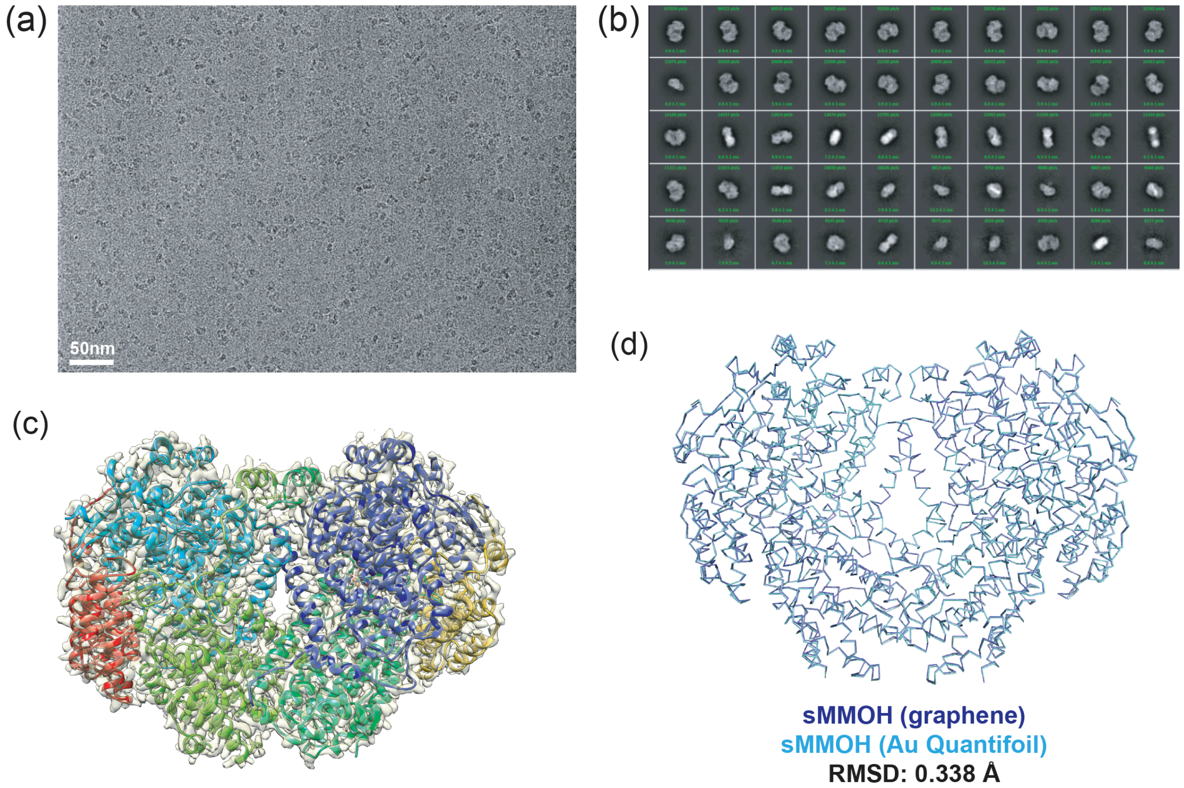


**Figure S4.** (a) Representative microscopic image of *M. caps* sMMOH in the Au Quantifoil grid (300 KeV). (b) Top 50 2D classes selected from 200 classes of sMMOH in the Au Quantifoil grid. (c) 2.9 Å resolution cryo-EM map of sMMOH in the Au Quantifoil grid. Two MMOHα subunits are colored in blue/cyan, MMOHβ in green/light green, and MMOHγ in yellow/red. (d) Structural overlay of sMMOH cryo-EM structures determined using Au Quantifoil (cyan) and graphene (blue) grids. The Cα R.M.S. difference in between is 0.338 Å.

### Preferred Particle Orientation

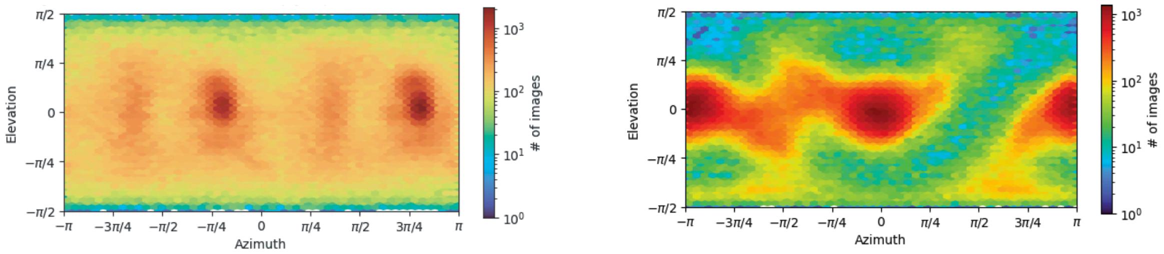


**Au Quantifoil (Final 479 k particles)**

**Graphene (Final 325 k particles)**

**Figure S5.** The preferred particle orientation of sMMOH determined by Au Quantifoil (479 k particles) and graphene (325K particles) grids. The images were imported from cryoSPARC.

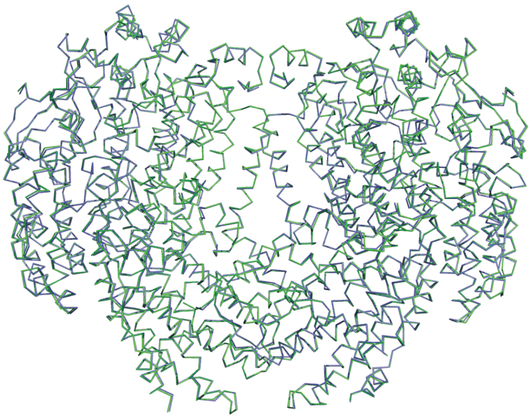


**Figure S6.** The structural overlay of the x-ray crystal structure (green; PDB ID: 1MTY) and cryo-EM structure (blue; graphene grid) of sMMOH. The Cα R.M.S. difference of two structures was 0.351 Å.

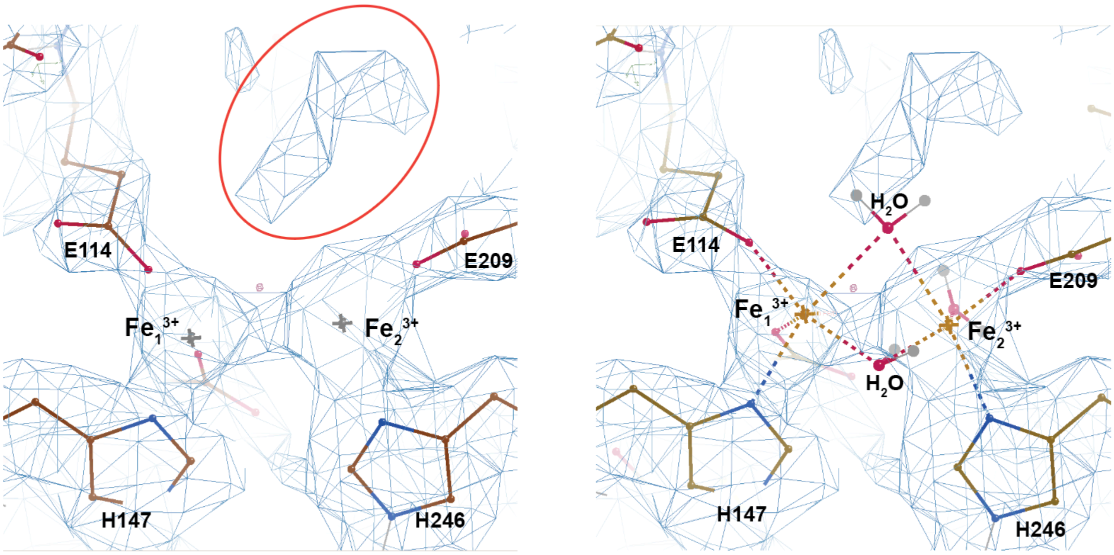


**Figure S7.** Cryo-EM map of sMMOH at the di-iron center (graphene grid) was overlaid with the cryo-EM structure (left) and x-ray structure (right; PDB ID: 1MTY) of sMMOH. The extra unidentified cryo-EM density on top of the di-iron center is displayed and marked by red circle.
